# CPSF3 inhibition blocks pancreatic cancer cell proliferation through disruption of core histone processing

**DOI:** 10.1101/2022.05.09.491230

**Authors:** Abdulrahman. A. Alahmari, Aditi H. Chaubey, Arwen A. Tisdale, Carla D. Schwarz, Abigail C. Cornwell, Kathryn E. Maraszek, Emily J. Paterson, Minsuh Kim, Swati Venkat, Eduardo Cortes Gomez, Jianmin Wang, Katerina V. Gurova, Michael E. Feigin

**Author notes:** Correspondence may be addressed to: MEF, Twitter: @TheFeiginLab.

## Abstract

Pancreatic ductal adenocarcinoma (PDAC) is a lethal disease with limited effective treatment options. This potentiates the importance of uncovering novel drug targets. We have discovered global dysregulation of the gene regulatory process alternative polyadenylation (APA) in PDAC. APA is a pre-mRNA processing mechanism that generates mRNAs with distinct 3’ ends, impacting gene expression and protein function. We revealed that APA dysregulation in PDAC drives oncogenic signatures and predicts poor patient outcome. As APA directs widespread gene expression dysregulation across the PDAC patient population, we hypothesized that inhibition of APA has therapeutic potential. APA is controlled by a complex of proteins, including cleavage and polyadenylation specificity factor 3 (CPSF3). CPSF3 is the endonuclease catalyzing mRNA cleavage, and a potentially druggable target. We now find that *CPSF3* is highly expressed and associated with poor prognosis in PDAC patients. CPSF3 knockdown decreases PDAC proliferation and clonogenicity *in vitro* and tumor growth *in vivo*. We demonstrate that *CPSF3* knockdown induces widespread APA alterations of oncogenes and tumor suppressors, and determine the contribution of one of these events to CPSF3-induced cell proliferation phenotype. Furthermore, we find that PDAC, but not non-transformed pancreatic cells, are sensitive to the CPSF3 small molecule inhibitor JTE-607. Mechanistically, JTE-607 impairs replication-dependent histone processing, disrupting nucleosome assembly and destabilizing chromatin structure. Finally, we determine that JTE-607 attenuates cell proliferation by arresting cells in early S-phase of the cell cycle. Altogether, we identify CPSF3 as a druggable target in PDAC and reveal novel mechanisms by which CPSF3 controls cancer cell growth.

**Significance:** This work identifies CPSF3 as a potential drug target in pancreatic ductal adenocarcinoma and reveals new mechanisms by which CPSF3 inhibition attenuates PDAC cell proliferation through modulating alternative polyadenylation and histone processing.

## Introduction

Pancreatic ductal adenocarcinoma (PDAC) is the third leading cause of cancer deaths with a five-year survival rate of 11%, due in part to the lack of effective treatment options (1). PDAC is primarily driven by mutations in the oncogene *KRAS* and several tumor suppressors, including *TP53*, *CDKN2A* and *SMAD4* (2). However, as clinically effective modulators of activity of these proteins are not currently available, identification of novel targets amenable to small molecule inhibition is a critical undertaking. Recently, large-scale RNA sequencing efforts of PDAC tumors have revealed widespread dysregulation of oncogenic gene expression, allowing the characterization of several PDAC subtypes and phenotypic states (3–6). These gene expression changes are critical for driving tumor phenotypes, including metastatic progression (7–11). While these gene expression changes have been extensively catalogued, the mechanisms underlying this transcriptional heterogeneity remain largely unknown (12). We propose that targeting these drivers of dysregulated gene expression represents an opportunity to reverse widespread oncogenic activity in transformed cells.

One such gene regulatory process that has been implicated in cancer is alternative polyadenylation, or APA (13–15). APA is a co-transcriptional mRNA processing mechanism that generates distinct transcript isoforms with different 3’ untranslated region (UTR) lengths, ultimately affecting mRNA stability, localization and translation (13). Recently, we identified widespread APA alterations in PDAC patients that are associated with functional changes in both gene and protein expression of growth-promoting genes (16). We revealed APA as a new mechanism that regulates PDAC gene expression, identified patterns of APA associated with poor patient outcome, and uncovered novel APA-regulated therapeutic targets. As APA is widely dysregulated across the PDAC patient population, and drives known pro-tumorigenic pathways, we propose that APA inhibition represents a promising therapeutic approach.

Cleavage and polyadenylation specificity factor 3 (CPSF3) is the endonuclease responsible for mRNA cleavage and is part of two distinct RNA processing complexes (17). One is the APA complex where CPSF3 cooperates with other APA factors to cleave the mRNA prior to the addition of the poly(A) tail. The second is the histone cleavage complex (HCC) where pre-mRNAs of replication-dependent core histones are cleaved by CPSF3 but not polyadenylated. Recently, CPSF3 was identified as the target of the small molecule JTE-607 (18, 19). JTE-607 is hydrolyzed into an active compound that directly interacts with the CPSF3 interfacial cavity (19). This interaction inhibits CPSF3 catalytic activity leading to accumulation of unprocessed newly synthesized pre-mRNAs. JTE-607 induces apoptosis of human acute myeloid leukemia (AML) and Ewing sarcoma cells *in vitro* and prolongs survival of tumor-bearing mice in xenograft models *in vivo* (20, 21). Notably, administration of JTE-607 in healthy volunteers demonstrated the safety of this compound in humans, with no severe adverse events reported (22). However, the role of CPSF3 and the effect of JTE-607 in epithelial cancers remains largely unknown.

Here, we show that knockdown and/or inhibition of CPSF3 attenuates PDAC cell proliferation *in vitro* and *in vivo*. We find that CPSF3 is highly expressed in PDAC patients and is a predictor of poor outcome. We conduct the first global analysis of CPSF3 loss in cancer, uncovering APA events correlating with expression alterations in tumor suppressors and oncogenes. Additionally, we find that small molecule inhibition of CPSF3 by JTE-607 selectively attenuates proliferation of PDAC cells but not non-transformed cells. Finally, we uncover a new mechanism by which JTE-607 attenuates cell proliferation, through disruption of replication-dependent histone mRNA processing, thus altering chromatin stability and dysregulating the cell cycle.

To our knowledge, our study is the first to pharmacologically target CPSF3 activity in epithelial cancers in general, and in PDAC specifically. We reveal that CPSF3 disruption blocks cell proliferation through multiple mechanisms, including APA-mediated gene expression alterations and disruption of proliferation-dependent histone mRNA processing. Furthermore, we provide the first connection between CPSF3 inhibition and chromatin stability. Overall, our findings uncover new functions of CPSF3 in cancer and nominate CPSF3 as a novel therapeutic target in PDAC.

## Materials and Methods

### Cell lines and *in vitro* culture

HEK293T, MiaPaCa2 and Panc1 were purchased from ATCC. Suit2 cells were obtained from Dr. David Tuveson (Cold Spring Harbor Laboratory). Human immortalized C7 CAFs and PancPat CAFs cells were obtained from Dr. Edna Cukierman (Fox Chase Cancer Center). Cells were cultured in DMEM (Corning, DMEM [+] 4.5 g/L glucose, L-glutamine, sodium pyruvate) supplemented with 10% fetal bovine serum (FBS) and 1% penicillin-streptomycin. Non-transformed pancreatic cell lines HPNE and HPDE were obtained from Dr. Ethan Abel (Roswell Park Comprehensive Cancer Center). HPNE cells were cultured in 75% DMEM+25% Medium M3 Base supplemented with 2mM L-glut, 1.5g/L sodium bicarbonate, 5% FBS, 10ng/mL hEGF and 5.5mM D-glucose. HPDE cells were cultured in Keratinocyte SFM (serum-free media) supplemented with 25mg BPE, 2.5μg EGF, 1X anti-anti and 50μg/mL Gentamicin. HeLa-TI cells were obtained from Dr. Katerina Gurova (Roswell Park Comprehensive Cancer Center) and were cultured in phenol red-free FluoroBrite DMEM complete Media (ThermoFisher). All cell lines were cultured at 37°C with 5% CO_2_ and tested negative for Mycoplasma.

### Generation of stable CPSF3 knockdown cells

Vectors expressing short-hairpin RNA (shRNA) targeting CPSF3 (sh1 Target sequence: GCTGAGATTGATCTCCTATTA; Clone ID: NM_016207.2-219s1c1, sh2 Target sequence: CCAGTGAATTTATTCGTGCTT; Clone ID: NM_016207.2-1240s1c1) were purchased from Sigma-Aldrich. Cells were infected with lentivirus harboring pLKO.1-shNTC (non-targeting control) and pLKO.1-shCPSF3. Polybrene was used to increase the efficacy of infection. After 72 hours, cells were selected with 2.5μg/ml puromycin. Knockdown was confirmed by qPCR and immunoblotting.

### RNA isolation and quantitative PCR

Cells were lysed with TRIzol reagent (Thermofisher Scientific; Cat # 15596026). RNA was isolated using Direct-zol RNA Miniprep Kit (Zymo Research; Cat # R2050). cDNA was synthesized using iScript cDNA Synthesis Kit (Bio-Rad; Cat# 1708891). qPCR was conducted with SYBR Green PCR primers (CPSF3; Unique Assay ID: qHsaCID0007422, ACTB; Unique Assay ID qHsaCED0036269, HIST1H3B; Unique Assay ID: qHsaCED0007746, HIST1H2BC; Unique Assay ID: qHsaCED0007746, FHL1; Unique Assay ID: qHsaCED0038537) mixed with iTaq Universal SYBR Green Supermix (Bio-Rad; Cat# 1725120) and run on CFX connect systems (Bio-Rad). Data were analyzed in Microsoft Excel and graphed using GraphPad Prism (v 9.3.0).

### RNA-sequencing and APA analysis

Cells were trypsinized, washed with 1X PBS and sent frozen (-80°C) for RNA sequencing (RNA-seq). 500ng total RNA was used to prepare the sequencing libraries using KAPA RNA HyperPrep Kit with RiboErase (HMR) (Roche Sequencing Solutions) following manufacturer’s protocol. Briefly, ribosomal RNA (rRNA) was depleted from total RNA and DNase-digested to remove gDNA contamination. RNA was purified, fragmented and first strand cDNA was synthesized using random primers. cDNA:RNA hybrids were converted into double-stranded cDNA (dscDNA) using dUTP incorporation. Adapters were added to the 3’ ends, ligated to library insert fragments and the library amplified in a strand-specific manner. Data were analyzed by the Bioinformatics Shared Resource (Roswell Park Comprehensive Cancer Center). APA analysis was done using Dynamic Analysis of Alternative Polyadenylation from RNA-seq (DaPars) algorithm (23). Exact steps followed for this analysis were previously described in (16).

### Immunoblotting

Samples were lysed using RIPA lysis buffer (50mM Tris. HCl pH 7.5, 150mM NaCl, 5mM EDTA pH 8, 1% Triton X-100, 0.5% NP-40) in the presence of 10ug/ml protease inhibitors (Aprotinin, Leupeptin, PMSF), boiled at 95°C for 5min and resolved by SDS-PAGE. Proteins were transferred to nitrocellulose membranes (0.2 µm, Bio-Rad, Cat. # 1620112) at a constant voltage of 100V for 70 minutes at 4°C using Mini Trans-Blot® Cell (Bio-Rad). Membranes were blocked in TBST (Tris-buffered saline (TBS) with 0.1% v/v TWEEN-20; Sigma Aldrich) and 5% w/v nonfat dry milk (Blotting-Grade Blocker #1706404, Bio-Rad). Primary antibodies were diluted in 3% BSA in TBST and incubated overnight at 4°C (mouse monoclonal CPSF3 antibody, Abcepta, AT1610a; rabbit polyclonal Histone H3 antibody, Cell Signaling Technology, 9715S; rabbit polyclonal Histone H2B antibody, Cell Signaling Technology, 8135S; mouse monoclonal GAPDH antibody, Proteintech, 60004-1-Ig; rabbit polyclonal FHL1 antibody, Proteintech, 10991-1-AP). Membranes were incubated with horseradish peroxidase-conjugated secondary antibodies at 1:2,000 dilution (Donkey anti-rabbit; Fisher Scientific; Catalog number: 45-000-682, or Goat anti-mouse; Sigma-Aldrich; Catalog number: A4416) for 1 hour at room temperature. Pierce ECL Western Blotting Substrate (Thermo Scientific, Catalog number: 32106) and Supersignal West Femto Maximum Sensitivity substrate (Thermo Scientific, Catalog number: 34094) were used for chemiluminescent detection. Signals were visualized and imaged using the ChemiDoc XRS+ System and Image Lab Software (Bio-Rad).

### Proliferation and clonogenicity assays

For proliferation experiments, cells were seeded at a density of 250 cells/well (MiaPaCa2 and Panc1 cells) or 1000 cells/well (HPNE cells) into a white 96-well plate in triplicate. Cell proliferation was measured using CellTiter-Glo Luminescent Cell Viability Assay Kit (Promega) at days 0, 2, 4 and 6. For clonogenicity assays, cells were seeded at a density of 500 cells/well into a 6-well plate in triplicate. After 11 days, cells were fixed with 4% formalin, stained with 0.2% Crystal Violet and images were obtained for analysis. Colony area was measured using ImageJ software. Data were normalized to control data points.

### Cell cycle analysis

Cells were trypsinized and resuspended in 1X PBS, then fixed with ice-cold 70% ethanol for 1 hour at -20°C. Cells were then washed with cold 1X PBS and incubated with RNaseA (200μg/ml) at 37°C for 1 hour. Propidium iodide (40μg/ml) was then added, incubated for 1 hour in the dark and analyzed by FACS at 488nm. Data were analyzed by FCS express software (v7.06.0015).

### BrdU incorporation assay

Cells were cultured under optimum conditions and incubated with 50μM BrdU (5-Bromo-2′- deoxyuridine; Sigma-Aldrich; B5002) for 4 hours. Cells were then rinsed with 1X PBS, trypsinized, permeabilized in 70% ice cold ethanol with gentle vortexing and stored at -20°C overnight. Next, cells were pelleted and DNA was hydrolyzed by incubating with 500μl of 2N HCl, 0.5% Triton X-100 in 1X PBS, incubated for 30 minutes at room temperature and then neutralized by adding 1.5ml of 0.1 M sodium tetraborate (pH 8.5) for 2 minutes. Cells were then pelleted, washed once with 1% BSA in 1X PBS and resuspended in 50μl 0.5% Tween 20, 1% BSA in 1X PBS. Next, 10^6^ cells were incubated with 1μg Anti-BrdU-FITC (FITC anti-BrdU Antibody; BioLegend; Cat#364104) for 1 hour at room temperature. Cell pellets were washed once with 150µl 1% BSA in 1X PBS, resuspended in 500μL 1X PBS with RNaseA (200 μg/ml) and PI (40 μg/ml) and incubated at room temperature for 30 minutes in the dark. Cells were analyzed by flow cytometry immediately and a compensation step was performed. Data were analyzed by FCS Express software (v7.06.0015).

### 3’ RACE

3’ RACE was performed as previously described (16). Briefly, first strand cDNA was synthesized from the poly(A) tail using an adapter primer (5′-GACTCGAGTCGACATCGATTTTTTTTTTTTTTTTT-3′). Gene specific PCR amplification was then performed using a gene specific primer spanning the stop codon (FHL1: 5’-TCCACTGCAAAAAATGCTCCGTGA-3’) and an adapter-targeting primer (5′-GACTCGAGTCGACATCG-3′). The PCR products were run on a 1.2% agarose gel and visualized and imaged using the ChemiDoc XRS+ System and Image Lab Software (Bio-Rad).

### Xenograft experiments

Animal experiments were approved by the Roswell Park Institutional Animal Care and Use Committee. MiaPaCa2 cells infected with shNTC and sh1 CPSF3 constructs were trypsinized, washed with 1X PBS and counted. 5x10^5^ cells were resuspended in 50μl of 1X PBS/Matrigel (Corning Life Sciences, 356231) in a 1:1 ratio and injected subcutaneously into the flanks of 8-week old NOD/SCID/IL2Rγ^-/-^ (NSG) mice. When palpable, tumor volume was determined by caliper measurements obtained in 2 dimensions and calculated as width^2^ x length/2 twice a week. Mice were euthanized when the first tumor reached 1400 mm^3^, tumors were dissected, and tumor volumes were measured.

### JTE-607 studies

For dose-response measurements, cells were seeded at a density of 1000 cells per well in a 96- well white plate. The next day, JTE-607 was titrated over a range of concentrations using the Tecan D300e Digital Dispenser and cell viability was measured 72 hours post drug titration using a CellTiter-Glo Luminescent Cell Viability Assay Kit (Promega). For cell proliferation experiments, cells were seeded at a density of 250 cells per well in a 96-well white plate. DMSO control or JTE-607 was dispensed at varying concentrations and proliferation was measured using CellTiter-Glo Assay at days 0, 2, 4 and 6. For clonogenicity experiments, cells were seeded at a concentration of 500 cells per well and treated with different concentrations of JTE-607. Cells were allowed to grow over a period of 11-14 days after which they were fixed in 4% formalin, stained with 0.2% Crystal Violet and images were obtained for analysis. Colony area was measured using ImageJ software. Data were normalized to DMSO control data points.

### Micrococcal digestion

Micrococcal Nuclease (MNase) was performed as previously described (24). Briefly, cells were trypsinized, washed with 1X RSB buffer (10mM Tris HCL, pH7.6; 15mM NaCl; 1.5mM MgCl2) and pelleted at 1000rpm for 4 minutes at room temperature. Cell pellets were resuspended in 1X RSB buffer with 1% TritonX-100, homogenized with a loose pestle (5 strokes) and centrifuged for 5 minutes at 2000rpm at 4°C. Pellets were washed two times with 1ml of buffer A (10mM Tris HCL, pH7.6; 15mM NaCl; 60mM KCl; 0.34M Sucrose; 0.1% B-mercaptoethanol; 0.15mM Spermine; 0.5 mM Spermidine; 0.25mM PMSF) and nuclei were pelleted at 160g for 10 minutes at 4°C. Nuclei were resuspended in 1.5ml of buffer A supplemented with 1mM of CaCl_2_. Nuclear suspensions (500μl) were digested with 200U/ml Micrococcal nuclease (NEB, cat#MO247S) at 37°C at different time points. Digestion was inactivated by 15mM EDTA. 10%SDS and 1M NaCl were added to extract genomic DNA. DNA was run and visualized using TapeStation 4200 system (Genomic Shared Recourse, Roswell Park Comprehensive Cancer Center).

### Statistical analyses

Experimental findings were obtained from three independent experiments unless stated otherwise. Statistics were performed in GraphPad Prism 9. In general, P< 0.05 was considered statistically significant. All statistical methods and P-values are provided in the figure legends. Asterisks in graphs denote statistically significant differences as described in figure legends.

### Code availability

The code used to analyze the data is available at (https://github.com/feiginlab/CPSF3_PDAC).

## Results

### *CPSF3* is upregulated in human PDAC and predicts poor patient outcome

To determine the clinical significance of *CPSF3* expression in PDAC, we first analyzed gene expression data from the Clinical Proteomic Tumor Analysis Consortium (CPTAC) (25). *CPSF3* expression was significantly higher in PDAC tumors (n=135), as compared with non-tumor adjacent tissues (n=18) and normal pancreata (n=7) (Fig. 1A). Consistent with this finding, *CPSF3* expression was also significantly higher in the Pancreatic Adenocarcinoma (PAAD) dataset from The Cancer Genome Atlas (TCGA) (n=147) as compared to normal pancreata (n=165) from The Genotype-Tissue Expression (GTEx) project (Fig. 1B). Next, we sought to assess the relationship between *CPSF3* expression and PDAC patient outcome. Patients with high *CPSF3* expression had significantly worse overall survival than patients with low *CPSF3* expression (p=0.00164, hazard ratio 5.047 (1.842-13.827)). Specifically, patients in the top quartile of *CPSF3* expression had a median survival of 14.2 months, while those in the bottom quartile of *CPSF3* expression had a median survival of 33.5 months (Fig. 1C). We then sought to assess *CPSF3* expression status in our cell line models. In agreement with the clinical data, we found that *CPSF3* is upregulated in PDAC cell lines (MiaPaCa2, Suit2, Panc1) as compared to non-transformed pancreatic epithelial cells (HPNE, HPDE) by RT-qPCR and western blot (WB) (Fig 1D,E). Therefore, *CPSF3* is highly expressed in PDAC, high expression correlates with poor patient outcome, and our cell models are appropriate for mechanistic studies.

**Figure 1.**
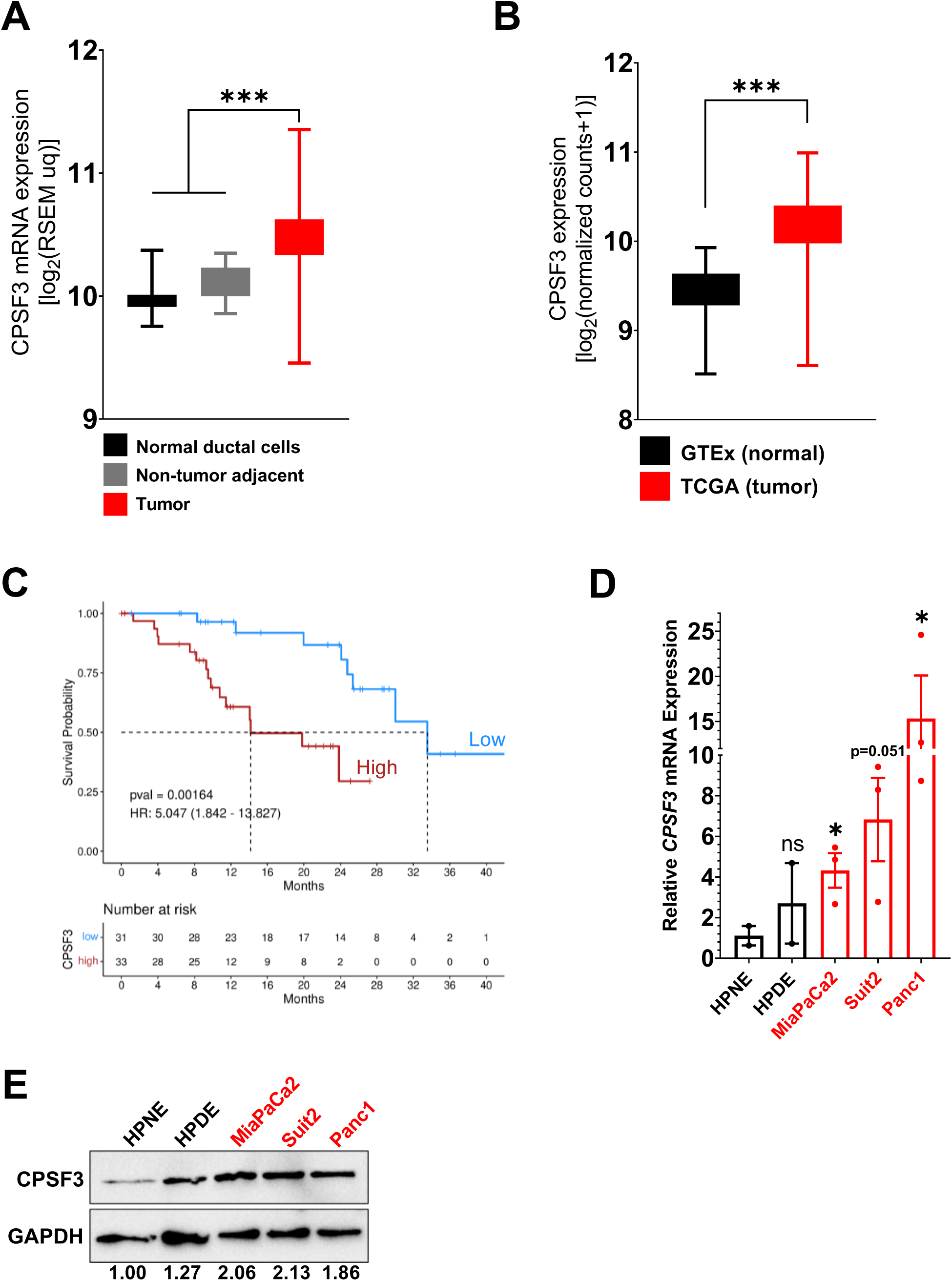
*CPSF3* is highly expressed in PDAC patients and predicts poor prognosis. **A,** *CPSF3* expression from CPTAC PDAC patient data. Whiskers indicate minimum and maximum data points. ***, *P*<0.0001, Ordinary one-way ANOVA with Tukey multiple comparisons test. **B,** *CPSF3* expression from PDAC patient data (TCGA) as compared to normal pancreas (GTEx). Whiskers indicate minimum and maximum data points. ***, *P*<0.0001, unpaired t test with Welch’s correction. **C,** Kaplan Meier survival curves of PDAC patients with high and low CPSF3 mRNA levels. Data were obtained from CPTAC database. **D,** Quantitative RT-PCR showing *CPSF3* mRNA expression levels in non-transformed pancreatic epithelial and PDAC cells. Data are shown as mean±SEM.*, *P* < 0.05, unpaired t test with Welch’s correction. **E,** Immunoblotting of CPSF3 in non-transformed pancreatic epithelial cells and PDAC cells. CPSF3 protein levels were normalized to its corresponding GAPDH levels. Fold change compares normalized data of different cell lines to HPNE.

### CPSF3 is required for PDAC cell proliferation and tumor growth

To define the functional role of *CPSF3* in PDAC we first took a genetic approach and generated stable *CPSF3* knockdown MiaPaCa2 and Panc1 cells. We used two different short hairpin RNAs (sh1 and sh2) targeting *CPSF3*, and a non-targeting control (shNTC). Successful knockdown of *CPSF3* was confirmed at the RNA and protein level by RT-qPCR and WB, respectively, with sh1 cells having the highest level of knockdown in both cell lines (Fig. 2A,B). We then examined the effect of *CPSF3* knockdown on cell proliferation and colony formation capability. *CPSF3* knockdown significantly attenuated proliferation as compared with shNTC controls in both MiaPaCa2 and Panc1 cells (Fig. 2C). *CPSF3* knockdown also significantly decreased colony formation (Fig. 2D,E). In both the proliferation and colony formation assays, and in both PDAC cell lines, sh1 CPSF3 had the strongest phenotype, consistent with higher levels of *CPSF3* knockdown.

**Figure 2.**
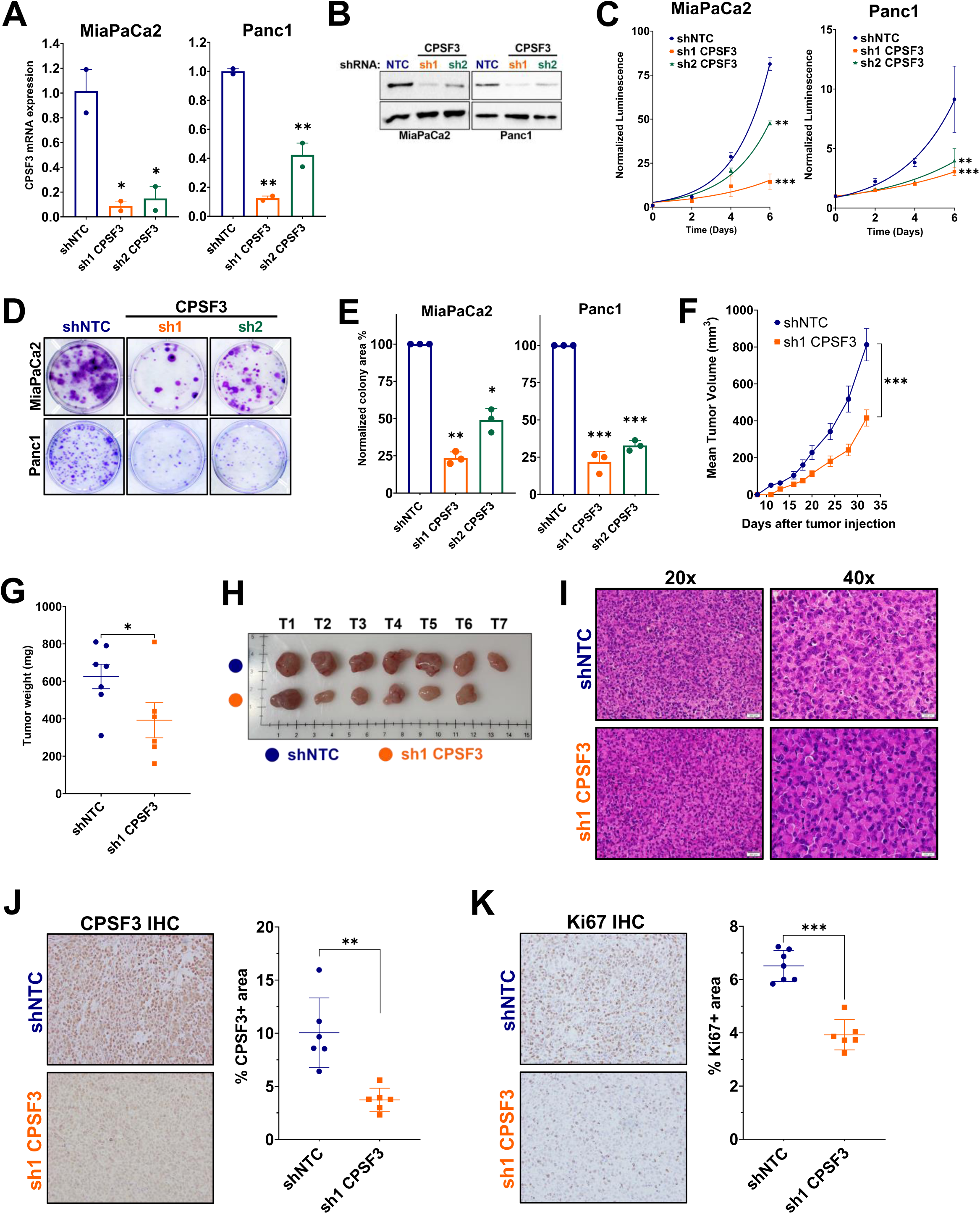
CPSF3 is required for PDAC cancer cell proliferation. **A,** mRNA expression of *CPSF3* in shNTC, sh1 CPSF3 and sh2 CPSF3 knockdown cells by qPCR. Graphs are representative of at least two independent experiments. Data are shown as mean±SEM of technical duplicates. **\***\*, *P* < 0.01, Ordinary one-way ANOVA with Dunnett’s multiple comparisons test. **B,** Immunoblotting of CPSF3 in shNTC, sh1 and sh2 CPSF3 knockdown cells. **C,** Proliferation rates at days 0, 2, 4 and 6 of shNTC, sh1 and sh2 CPSF3 knockdown cells. **\***\*, *P* < 0.01; **\*\***\*, *P* < 0.001; 2way ANOVA with Dunnett’s multiple comparisons test. **D,** Clonogenic growth assay of shNTC, sh1 and sh2 CPSF3 knockdown cells. **E,** Normalized colony area percentage of shNTC, sh1 and sh2 CPSF3 knockdown cells from **(D)**. *, *P* < 0.05; **\***\*, *P* < 0.01; **\*\***\*, *P* < 0.001; Ordinary one-way ANOVA with Dunnett’s multiple comparisons test. **F,** Volume of CPSF3-knockdown and control MiaPaCa2 tumors. ***, *P* < 0.001, 2way ANOVA. **G,** Endpoint tumor weight. *, *P* < 0.05, unpaired t test with Welch’s correction. **H,** Gross images of shNTC (n=7) and shCPSF3 (n=6) dissected tumors. **I,** Hematoxylin and Eosin (H&E) staining of xenograft tumors. **J** and **K,** IHC for CPSF3 and Ki67, respectively. Box and whisker plots indicate the percentage of CPSF3- and Ki67-positive areas in the tumors. **\***\*, *P* < 0.01; **\*\***\*, *P* < 0.001; unpaired t test.

Next, we sought to determine the requirement for CPSF3 in PDAC tumor growth *in vivo*. We implanted MiaPaCa2 cells (either shNTC or sh1 CPSF3, 5x10^5^ per mouse) subcutaneously into the flanks of NOD/SCID/IL2Rγ^-/-^ (NSG) mice. *CPSF3* knockdown tumors grew significantly slower, and weighed significantly less at endpoint, than shNTC tumors (Fig. 2F-H). No changes in tumor histopathology were noted by Hematoxylin and Eosin (H&E) staining (Fig. 2I). Immunohistochemical (IHC) analysis revealed that CPSF3 knockdown was maintained *in vivo* (Fig. 2J). Finally, IHC for Ki67 revealed a significant decrease in proliferation in *CPSF3* knockdown tumors as compared with shNTC controls (Fig. 2K). Overall, these data support the requirement for CPSF3 in PDAC cell proliferation and tumor growth.

### CPSF3 knockdown dysregulates global gene expression in PDAC cells

CPSF3 is an integral component of the polyadenylation complex; therefore, we hypothesized that *CPSF3* knockdown would dysregulate APA, leading to global changes in mRNA expression. To test this hypothesis, we subjected sh1 CPSF3 and shNTC Panc1 cells to RNA-sequencing (RNA-seq). Differential expression analysis revealed 376 genes significantly upregulated and 98 genes significantly downregulated (FDR<0.05; fold change >1.5) upon *CPSF3* knockdown. To uncover significantly altered changes in 3’-UTR length upon *CPSF3* knockdown, we applied the DaPars algorithm (23). DaPars identifies APA changes from standard RNA-seq data by generating a Percentage Distal Usage Index (PDUI) score for each gene based on the relative abundances of 3′-UTR long and short forms. A negative PDUI indicates a shortening event, while a positive PDUI indicates 3’-UTR lengthening. In accordance with our hypothesis, loss of *CPSF3* resulted in global APA dysregulation, with 402 genes having significantly shorter 3’-UTRs, and 292 genes having significantly longer 3’-UTRs in *CPSF3* knockdown cells (-0.1>PDUI>0.1; P<0.05) (Fig. 3A). APA-mediated alterations in 3’-UTR length impact mRNA stability and thus, gene expression (13,26– 28). To determine which CPSF3-mediated APA events might be driving individual changes in gene expression, we looked for genes that were significantly altered in both the DaPars and gene expression analyses (Fig. 3B). Intriguingly, this list included multiple downregulated oncogenes, including SMAD Family Member 6 (*SMAD6)* and Mitogen-Activated Protein Kinase Kinase 6 (*MAP2K6)*, and upregulated tumor suppressors, including Four And A Half LIM Domains 1 (*FHL1)* and CKLF-Like MARVEL Transmembrane Domain Containing 3 (*CMTM3)*. Therefore, these data support a role for *CPSF3* in controlling cell proliferation via APA-mediated dysregulation of a suite of cancer-associated genes.

**Figure 3.**
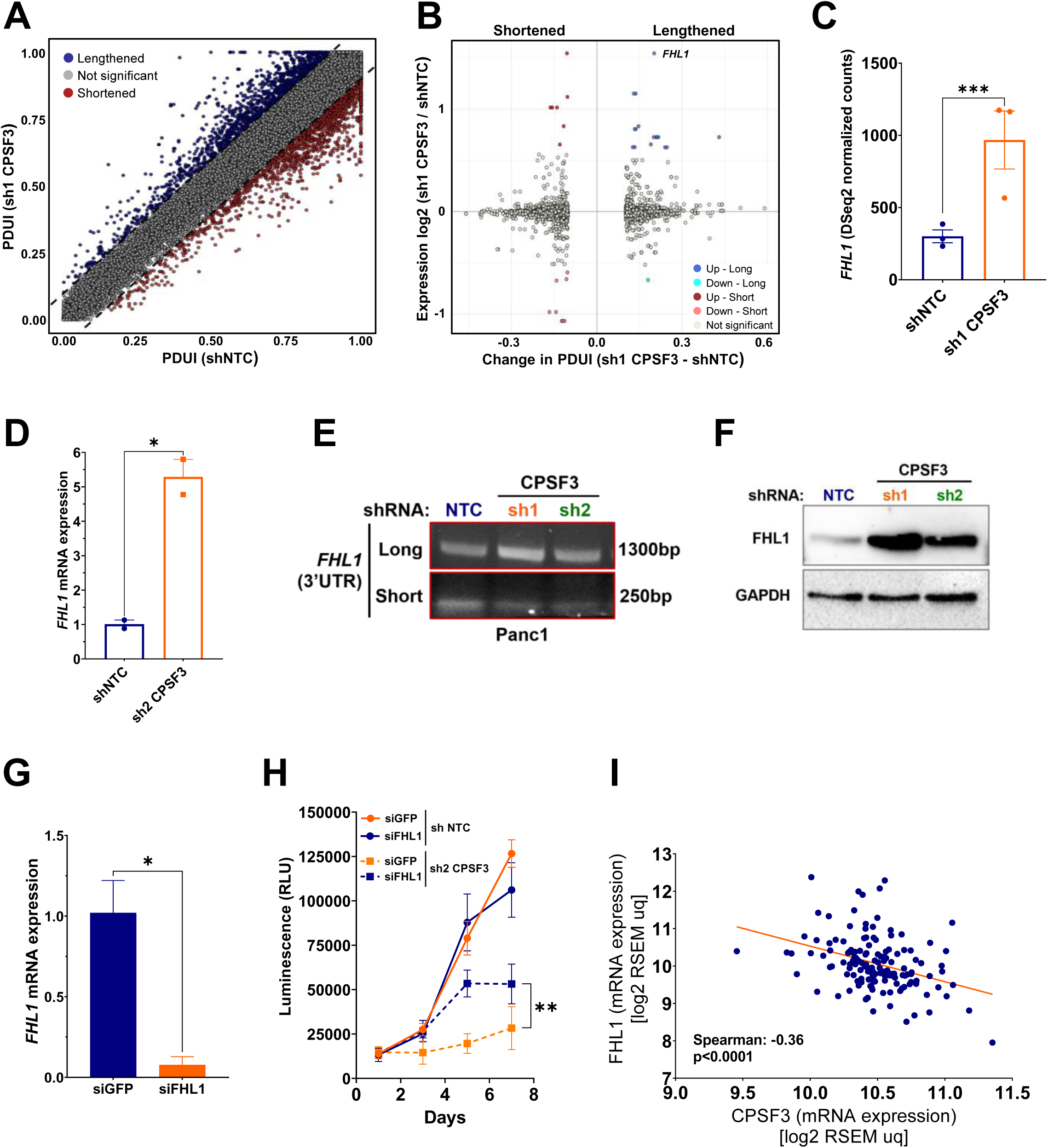
CPSF3 knockdown drives global APA dysregulation in PDAC. **A,** PDUI score of each gene in shNTC and shCPSF3 cells. Dashed lines represent 0.1 cutoffs. Blue dots represent 3′-UTR-lengthened genes, and red dots represent 3′-UTR-shortened genes. **B,** Genes showing lengthening (right) or shortening (left) events (-0.1> PDUI > 0.1; P<0.05) and are differentially expressed (FDR<0.05; fold change >1.5) as color coded. Up=upregulated gene expression, Down=downregulated gene expression. **C,** *FHL1* DSeq2 normalized counts. ***, *P* < 0.0001. **D**, *FHL1* mRNA levels in *CPSF3* knockdown MiaPaCa2 cells. *, *P* < 0.05, unpaired t test. **E,** 3’- RACE of FHL1 in shNTC, sh1 and sh2 CPSF3 Panc1 cells. Approximate length of the 3′-UTR form is denoted beside each band. **F,** Western blot of FHL1 protein levels in shNTC, sh1 and sh2 CPSF3 Panc1 cells. **G,** FHL1 mRNA levels in sh2 CPSF3 Panc1 cells transfected with siRNA against FHL1 (siFHL1). *, *P* < 0.05, unpaired t test. **H,** Proliferation assay of siFHL1 in *CPSF3* knockdown and control Panc1 cells. **, *P* < 0.01; 2way ANOVA with Tukey multiple comparisons test. **I,** Correlation of *FHL1* with *CPSF3* mRNA levels from the CPTAC database. Spearman’s correlation= -0.36. *P* < 0.0001.

We next sought to determine if any of the APA-regulated oncogenes or tumor suppressors were directly responsible for *CPSF3* knockdown-mediated attenuation of cell proliferation. We focused on *FHL1*, as it has been reported to possess tumor suppressor activity in non-PDAC cancers (29–32), but has no known roles in PDAC biology. FHL1 interacts with transcription factors and signaling proteins, thus modulating gene transcription and signaling pathways (29, 33). FHL1 suppresses tumor growth through several mechanisms, including interaction with tumor-regulating estrogen receptors and SMAD family proteins, and reduction of PI3K/AKT signaling (29,34,35). The *FHL1* 3’-UTR was significantly lengthened (P=0.029), and *FHL1* gene expression was significantly enriched, in *CPSF3* knockdown cells, as compared with shNTC controls (Fig. 3B,C). *FHL1* was also significantly overexpressed in *CPSF3* knockdown MiaPaCa2 cells, as compared with shNTC controls (Fig. 3D). We validated the shift in APA patterns of *FHL1* by 3′ RACE (rapid amplification of 3′ ends), revealing an increase in the long 3’-UTR form of *FHL1* in *CPSF3* knockdown cells, and a concomitant decrease in the abundance of the short 3’-UTR form (Fig. 3E). As APA can impact mRNA stability and translation, we determined FHL1 protein expression in shNTC and *CPSF3* knockdown cells by WB (Fig. 3F). We found that FHL1 protein levels were much higher in *CPSF3* knockdown cells. We then asked whether the CPSF3- mediated upregulation of FHL1 was responsible for the effect of *CPSF3* knockdown on PDAC cell growth. We transiently knocked down *FHL1* using siRNA in both shNTC and *CPSF3* knockdown cells and assessed the effect on proliferation. Knockdown of *FHL1* was greater than 90% effective (Fig. 3G). Knockdown of *FHL1* in shNTC cells had no effect on cell viability over the 7-day time course of the experiment (Fig. 3H). However, knockdown of *FHL1* significantly rescued the *CPSF3* knockdown-induced proliferation phenotype. Therefore, *CPSF3* knockdown reduces PDAC cell viability at least in part through upregulation of *FHL1*. Finally, we sought to determine if there was a correlation between *CPSF3* and *FHL1* gene expression levels in PDAC patients. We observed a significant negative correlation (R=-0.36, P<0.0001) between *CPSF3* and *FHL1* levels within the CPTAC database (Fig. 3I), supporting our conclusion that *FHL1* levels are mediated by *CPSF3* expression in PDAC.

### PDAC cells are sensitive to chemical inhibition of CPSF3

CPSF3 was recently identified as the target for the small molecule JTE-607. JTE-607 is a prodrug that, when metabolized by the ester hydrolyzing enzyme carboxylesterase 1 (CES1), binds to CPSF3 and inhibits its catalytic activity, impairing the processing of newly synthesized mRNAs (19). As genetic depletion of *CPSF3* attenuated PDAC cell proliferation (Fig. 2), we hypothesized that pharmacologic inhibition of CPSF3 with JTE-607 could represent a novel therapeutic approach in PDAC. We therefore examined the sensitivity of multiple human pancreatic cell lines, both non-transformed and PDAC, to JTE-607 in a 72-hour dose-response cell viability assay. Non-transformed pancreatic epithelial cells (HPNE, IC50=130.4μM; HPDE, IC50=60.11μM) and human cancer associated fibroblast cell lines (C7 CAF, IC50=70.04μM; PancPat CAFs, IC50=114.2 μM) were not sensitive to JTE-607 (Fig. 4A-C). In contrast, human PDAC cell lines displayed a range of sensitivities to JTE-607, with Panc1 cells being the most sensitive (IC50=2.163μM) (Fig. 4A,C). Next, we determined the effect of JTE-607 on cell proliferation by treating cells with increasing concentrations of JTE-607 and assessing cell viability in a time-dependent fashion (Fig. 4D,E). JTE-607 had no effect on proliferation in non-transformed HPNE cells (Fig. 4D). However, proliferation of MiaPaCa2 and Panc1 PDAC cells was significantly attenuated by JTE-607, in a dose-dependent manner (Fig. 4E). Finally, we tested the effect of JTE-607 on colony formation in PDAC cell lines. JTE-607 significantly decreased colony formation in all PDAC cell lines tested (Fig. 4F,G). Therefore, JTE-607 selectively attenuates proliferation of PDAC cells over non-transformed pancreatic cells.

**Figure 4.**
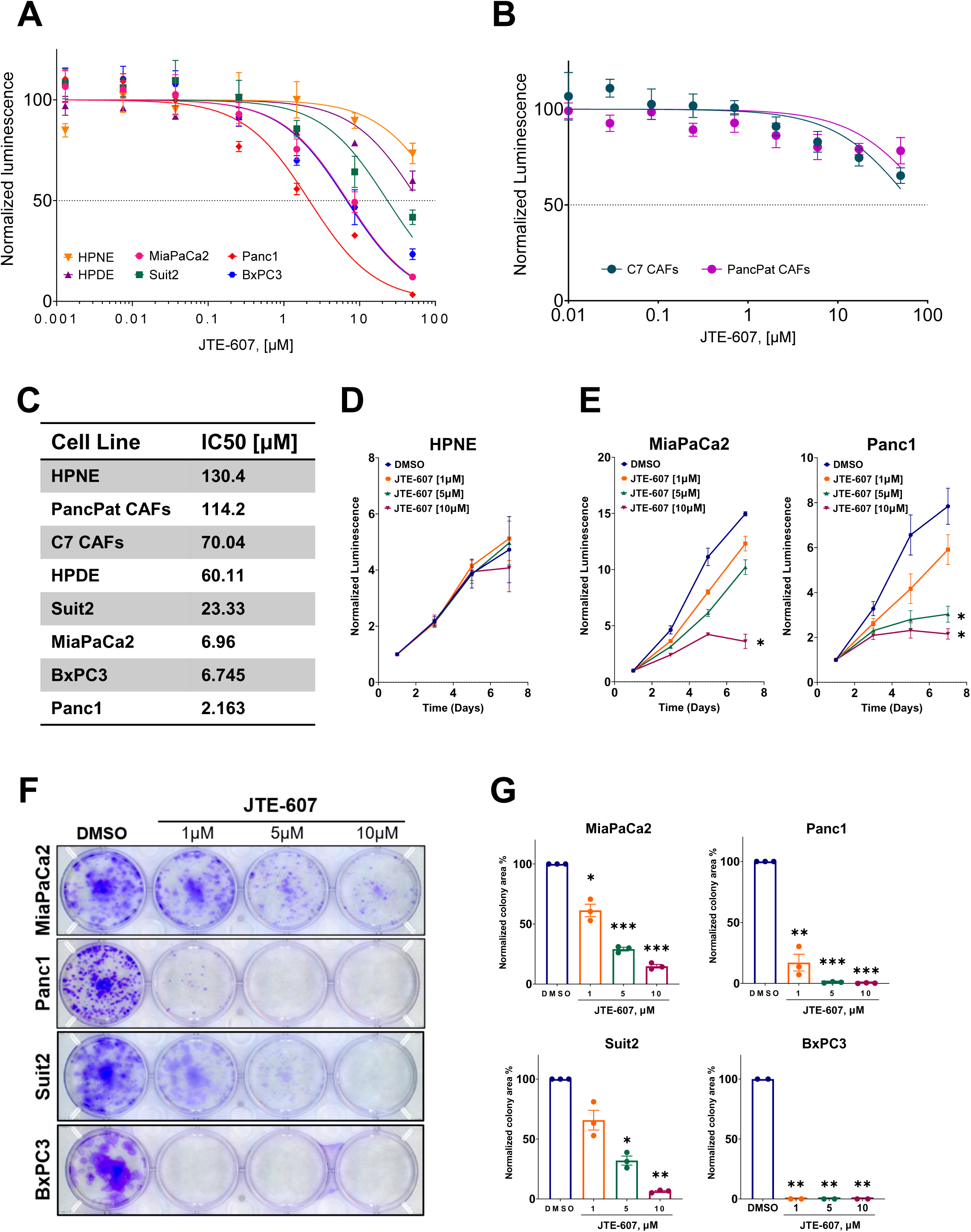
PDAC cell lines are sensitive to CPSF3 inhibition by JTE-607. **A,** IC50 of JTE-607 on non-transformed and PDAC cell lines. **B,** IC50 of JTE-607 on human fibroblast C7 and PancPat CAFs. **C,** IC50 values. **D and E,** Proliferation rates at days 0, 2, 4 and 6 of non-transformed and PDAC cell lines after treatment with escalating concentrations of JTE-607. *, *P* < 0.05; 2way ANOVA with Dunnett’s multiple comparisons test. Data are shown as mean±SEM. **F,** Clonogenic growth assay of PDAC cell lines after treatment with increasing concentration of JTE-607. **G,** Normalized colony area percentage of PDAC cell lines from **(F)**. *, *P* < 0.01; **\***\*, *P* < 0.001; **\*\***\*, *P* < 0.0001; Ordinary one-way ANOVA with Dunnett’s multiple comparisons test. Data are shown as mean±SEM.

### JTE-607 inhibits expression of replication-dependent histones

To determine the gene regulatory alterations underlying the ability of JTE-607 to attenuate PDAC cell proliferation, we performed RNA-seq on Panc1 cells treated with DMSO or JTE-607 for 24 hours. Differential gene expression and alterations in 3’-UTR length were analyzed as described above. Differential expression analysis revealed 1270 genes significantly upregulated and 646 genes significantly downregulated (FDR<0.05; fold change >1.5) upon JTE-607 treatment. We predicted that inhibition of CPSF3 would result in a global shift towards lengthened 3’-UTRs. In accordance with our hypothesis, JTE-607 resulted in global APA dysregulation, with 1242 genes having significantly longer 3’-UTRs, and 429 genes having significantly shorter 3’-UTRs in JTE-607 treated cells (-0.1>PDUI>0.1; P<0.05) (Fig. 5A). To determine which JTE-607-mediated APA events might be driving individual changes in gene expression, we looked for genes that were significantly altered in both 3’-UTR length and gene expression (Fig. 5B). This revealed multiple downregulated oncogenes, including N-Acetyltransferase 10 (*NAT10*) and Casein Kinase 1 Delta (*CSNK1D*), and upregulated tumor suppressors, including Elongation Factor For RNA Polymerase II 2 (*ELL2*) and Cyclin Dependent Kinase Inhibitor 1A (*CDKN1A*). Next, we compared global APA alterations between the *CPSF3* knockdown and JTE-607 treatment conditions. This analysis revealed an overlap in APA altered genes upon CPSF3 knockdown and JTE-607 treatment (120 genes altered in both conditions; -0.1>PDUI>0.1; P<0.05), supporting the contention that JTE-607 suppresses CPSF3 activity (Fig. 5C). However, many APA events were distinct between the two conditions, suggesting that long term CPSF3 knockdown may be mechanistically distinct from short term CPSF3 inhibition.

**Figure 5.**
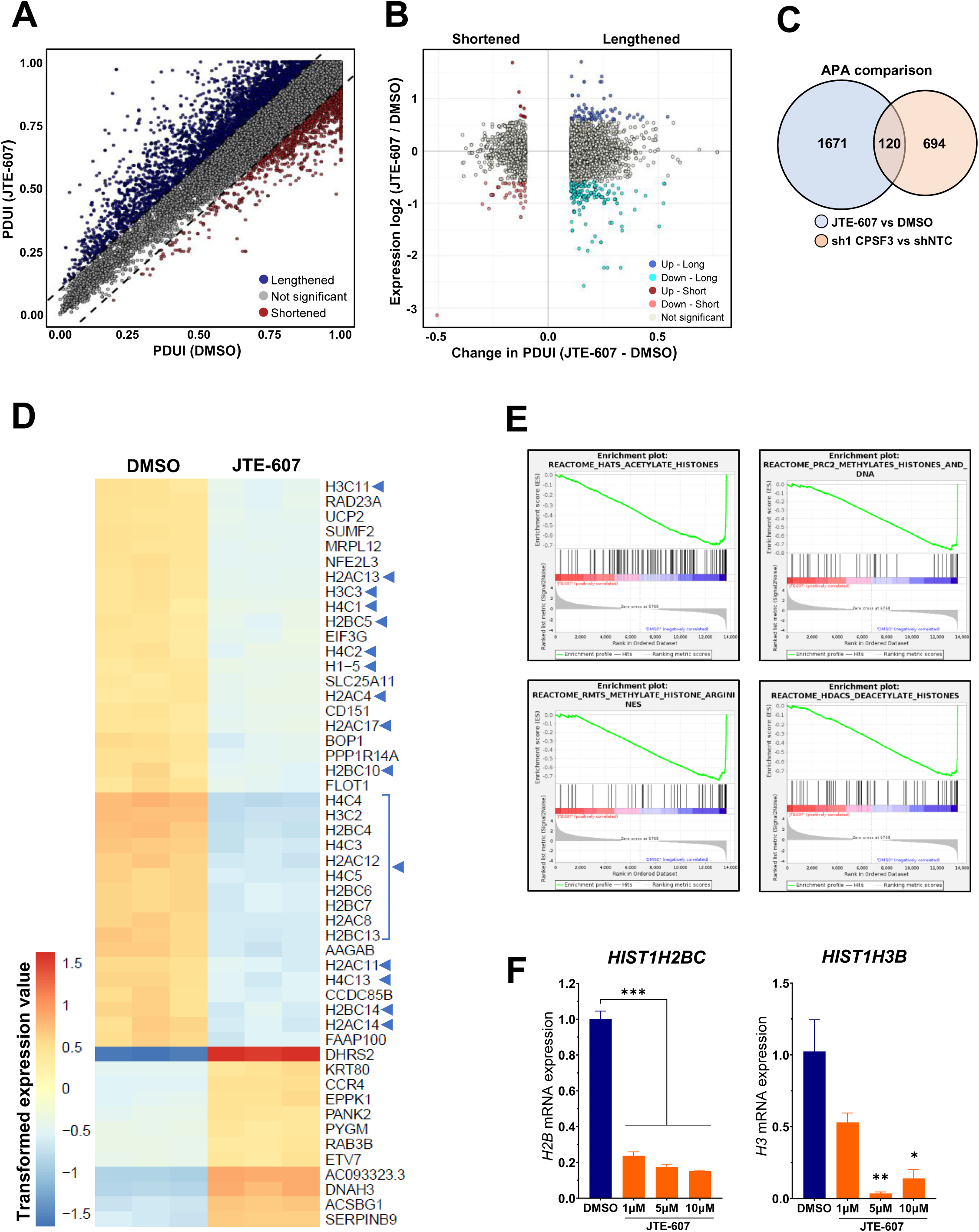
JTE-607 induces global APA dysregulation and decreases histone expression in PDAC cells. **A,** Changes in PDUI score (-0.1 > PDUI > 0.1; P<0.05) denoting 3′-UTR-shortened (red) and lengthened (blue) genes. **B,** Genes showing lengthening (right) or shortening (left) events (-0.1> PDUI > 0.1; P<0.05) and are differentially expressed (FDR<0.05; fold change >1.5) as color coded. Up=upregulated gene expression, Down=downregulated gene expression. **C,** Venn diagram showing overlapping genes with significant APA alterations between JTE-607- treated and *CPSF3* knockdown cells. **D,** Heatmap of differentially expressed genes in Panc1 cells treated with JTE-607. Expression is plotted as transformed expression value. Blue triangles denote replication-dependent histone genes. **E,** Gene set enrichment analysis (GSEA) of RNA-seq data from **(D)**. **F,** mRNA expression of *H2B* and *H3* in MiaPaCa2 cells treated with JTE-607. **\***, *P* < 0.05, **\*\***, *P* < 0.01, **\*\*\***, *P* < 0.001, Ordinary one-way ANOVA with Dunnett’s multiple comparisons test.

Next, we sought to determine how short term JTE-607 treatment attenuates PDAC proliferation. We were intrigued to find that numerous histone genes were significantly downregulated upon JTE-607 treatment in Panc1 cells (Fig. 5D). Gene set enrichment analysis (GSEA) demonstrated a dysregulation in many histone-related pathways, including histone methylation, acetylation and deacetylation (Fig. 5E). Interestingly, the majority of the differentially expressed histones were replication-dependent histones, including *HIST1H2AD*, *HIST1H2BJ*, *HIST1H3A* and *HIST1H4E*. We validated the JTE-607-induced decrease in replication-dependent histones in MiaPaCa2 cells (Fig 5F). Therefore, JTE-607 treatment decreases the expression of proliferation-dependent histones.

Replication-dependent histone genes are not polyadenylated and undergo pre-mRNA processing via the histone cleavage complex (HCC) (36). CPSF3 is the endonuclease component of the HCC (17,37,38); while *CPSF3* knockdown studies have demonstrated a role for CPSF3 in histone processing, the effect of inhibiting CPSF3 activity on histone mRNA processing has never been determined. We reasoned that lack of CPSF3-mediated cleavage activity would result in transcriptional read-through extending beyond the boundaries of the 3’-UTR. Indeed, we found that replication-dependent histones underwent transcriptional read-through upon JTE-607 treatment (Fig. 6A). Improperly processed mRNAs fail to be exported into the cytoplasm for translation, leading to decreased protein levels (17,39,40). Therefore, we examined replication-dependent histone protein levels upon JTE-607 treatment and found that JTE-607 reduced both H3 and H2B protein levels in a dose- and time-dependent fashion (Fig. 6B). In contrast, replication-independent histones did not undergo transcriptional read-through upon JTE-607 treatment, and did not decrease in gene expression levels (Fig. 6C,D). Finally, we sought to determine if there was a correlation between *CPSF3* and histone gene expression levels in PDAC patients. We calculated the Spearman’s correlation for *CPSF3* and 98 histone genes from the CPTAC database (Spearman = -0.15>R>0.15, P<0.05) (Fig. 6E-G). In accordance with our experimental findings, there were significantly more positive correlations (43 genes) between *CPSF3* and histone gene expression than negative correlations (3 genes) among replication-dependent histones (Fig. 5F). In contrast, there were few significant correlations between *CPSF3* and replication-independent histones (only 8 genes), and those significant alterations were equally positive and negative (Fig. 5G). Collectively, these results indicate that JTE-607 inhibits the function of CPSF3 in both the APA and histone mRNA processing complexes.

**Figure 6.**
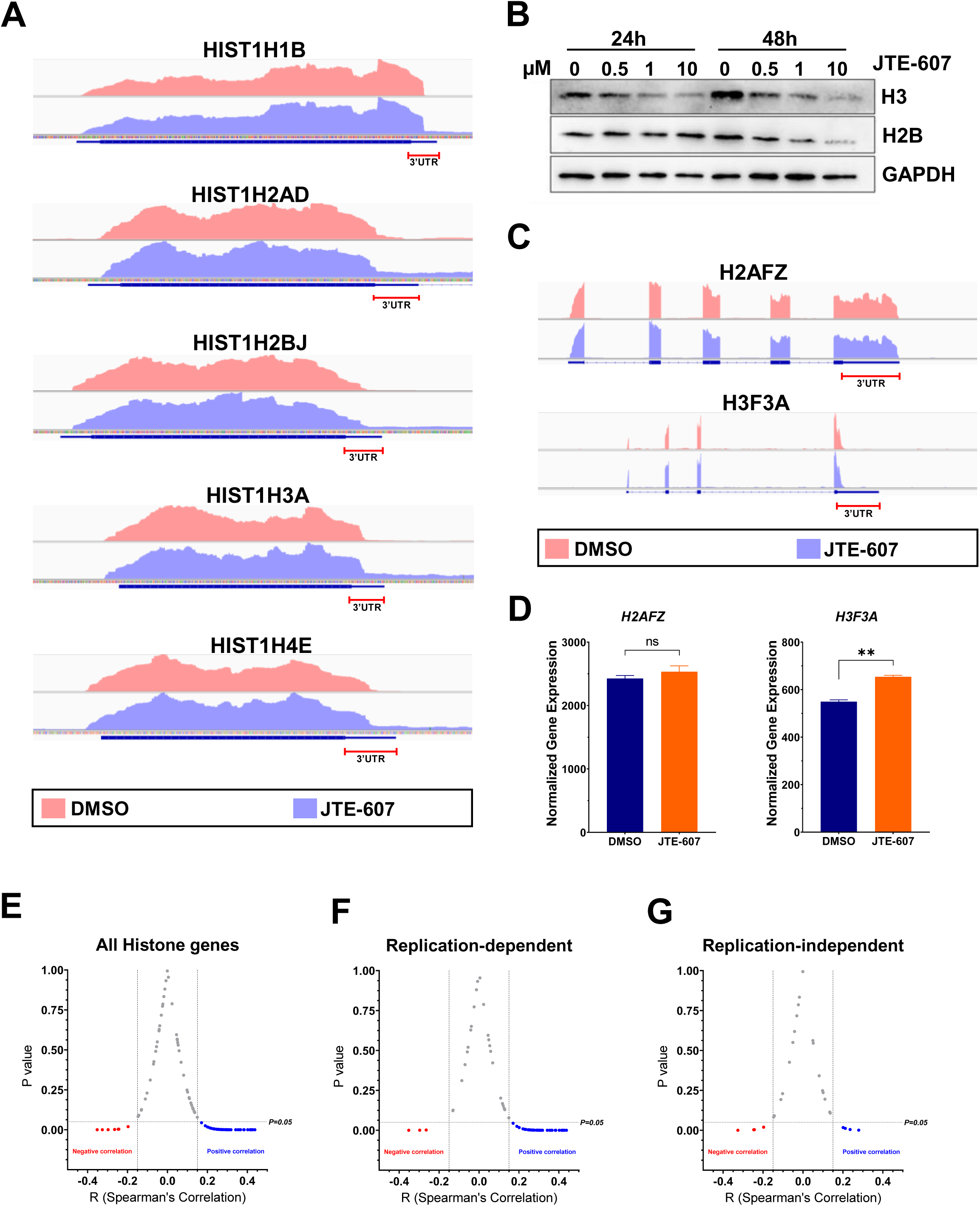
JTE-607 induces replication-dependent histone transcription read-through. **A,** IGV-generated density plots of replication-dependent histones highlighting the differences of 3′- UTR coverage between DMSO (red) and JTE-607 (blue) treated cells. **B,** Western blot of H2B and H3 protein levels in Panc1 cells treated with 0-10μM JTE-607 for 24 and 48hrs. **C,** IGV-generated density plots of replication-independent histones highlighting the differences of 3′-UTR coverage between DMSO (red) and JTE-607 (blue) treated cells. **D,** DSeq2 normalized counts of *H2AFZ* and *H3F3A* histone variants in Panc1 cells treated with JTE-607. **\*\***, *P* < 0.001. **E-G,** Volcano plots of Spearman’s correlation of CPSF3 and: **E,** all histone genes; **F,** replication-dependent histone genes; **G,** replication-independent histone genes. Each dot represents a histone gene. Blue and red dots denote positive and negative correlation, respectively. (Spearman = -0.15>R>0.15, P<0.05).

### JTE-607 destabilizes chromatin and blocks cell cycle progression

As replication-dependent histones are required for nucleosome assembly (36,41–43), we hypothesized that JTE-607 would dysregulate chromatin dynamics. First, we performed a Micrococcal Nuclease (MNase) assay to assess relative chromatin condensation. In this assay, protein-free DNA is digested by MNase, producing DNA fragmentation patterns that are indicators of whether chromatin is in a condensed or relaxed state. The chromatin destabilizing agent CBL0137 was used as a positive control (44). Panc1 cells treated with JTE-607 or CBL037 displayed rapid and complete chromatin digestion, as compared with DMSO-treated cells (Fig. 7A, quantification in 7B). To assess chromatin destabilization in a living cell, we utilized the HeLa-TI cell line model that has a silenced GFP reporter within a heterochromatic region of the genome. Treatment of these cells with chromatin destabilizing agents, including CBL0137, allows derepression of GFP silencing. Therefore, we monitored GFP expression in HeLa-TI cells upon JTE-607 treatment by both florescence microscopy and flow cytometry. Cells treated with JTE-607 induced GFP expression to levels comparable with CBL0137 in a dose- and time-dependent manner (Fig. 7C-E).

**Figure 7.**
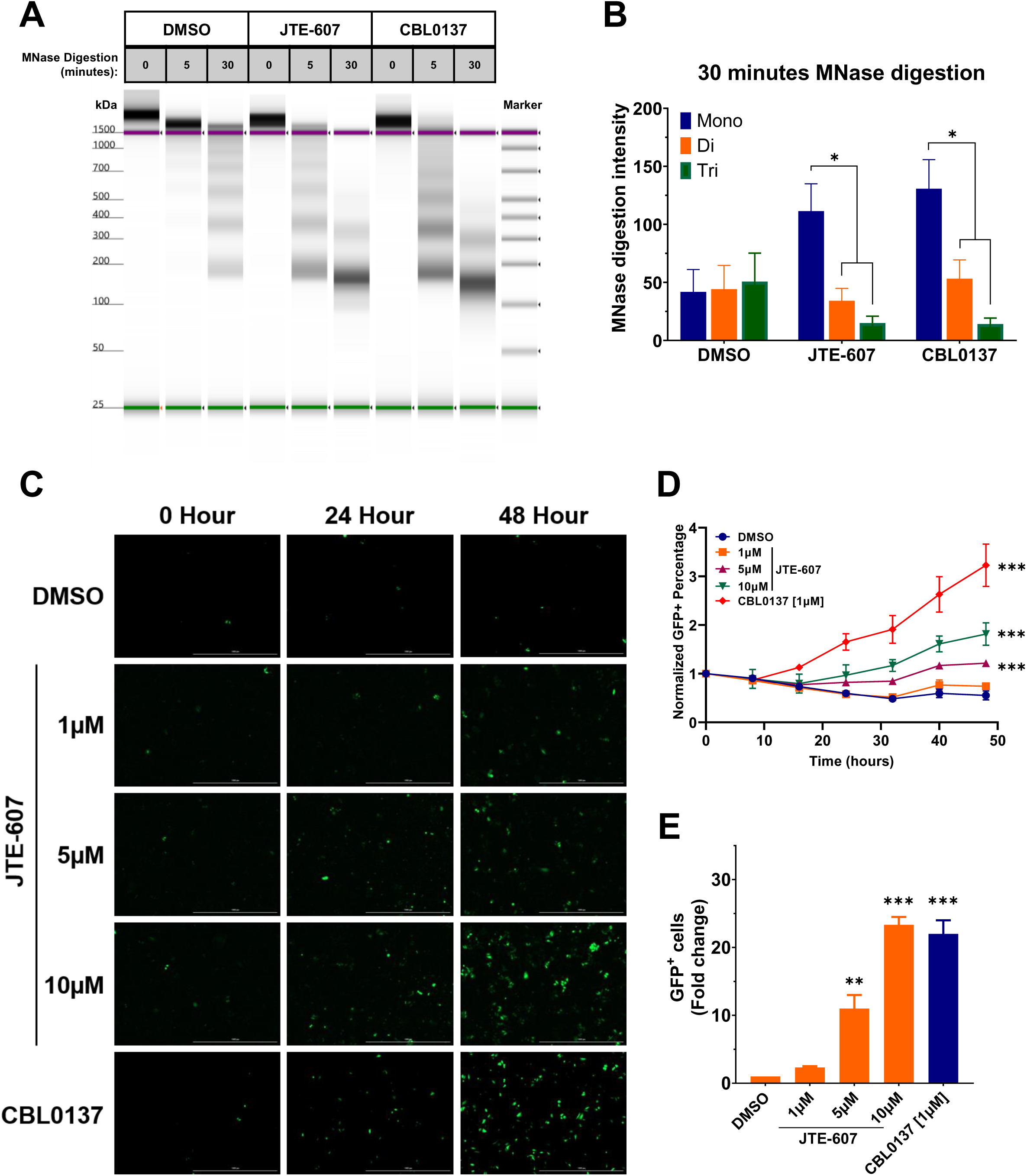
JTE-607 induces chromatin instability. **A,** Micrococcal Nuclease assay of Panc1 cells treated with JTE-607 or CBL0137. **B,** Quantification of nucleosome fragments after 30 minutes of MNase digestion. Data are shown as mean±SEM of two technical repeats. *, *P* < 0.05, unpaired t test with Welch’s correction. **C,** GFP+ HeLa-TI cells following JTE-607 or CBL0137 treatment. **D**, Fold change of GFP+ HeLa-TI from **(C).** ***, *P* < 0.0001; 2way ANOVA with Tukey’s multiple comparisons test. **E,** Flow cytometry analysis of GFP+ HeLa-TI cells following JTE-607 or CBL0137 treatment. Fold change is shown as mean±SEM of two independent experiments. **\*\***, *P* < 0.01, **\*\*\***, *P* < 0.0001, Ordinary one-way ANOVA with Tukey’s multiple comparisons test.

Finally, we sought to determine how JTE-607-mediated depletion of replication-dependent histones led to defects in cell viability. As replication-dependent histones are required for cell cycle progression, we assessed the effects of JTE-607 on cell cycle distribution. In non-transformed HPNE cells, JTE-607 had no impact on cell cycle distribution (Fig. 8A). In contrast, JTE-607 arrested Panc1 and MiaPaCa2 PDAC cells in S-phase of the cell cycle within 24 hours (Fig. 8B). To more specifically investigate the timing and extent of S-phase arrest, we examined BrdU incorporation upon JTE-607 treatment in a time-dependent manner (Fig. 8C). We found that JTE-607 arrests cells in early to mid S-phase of the cell cycle as early as 8 hours (Fig. 8C). Therefore, JTE-607 destabilizes chromatin and attenuates PDAC cell viability through S-phase cell cycle arrest.

**Figure 8.**
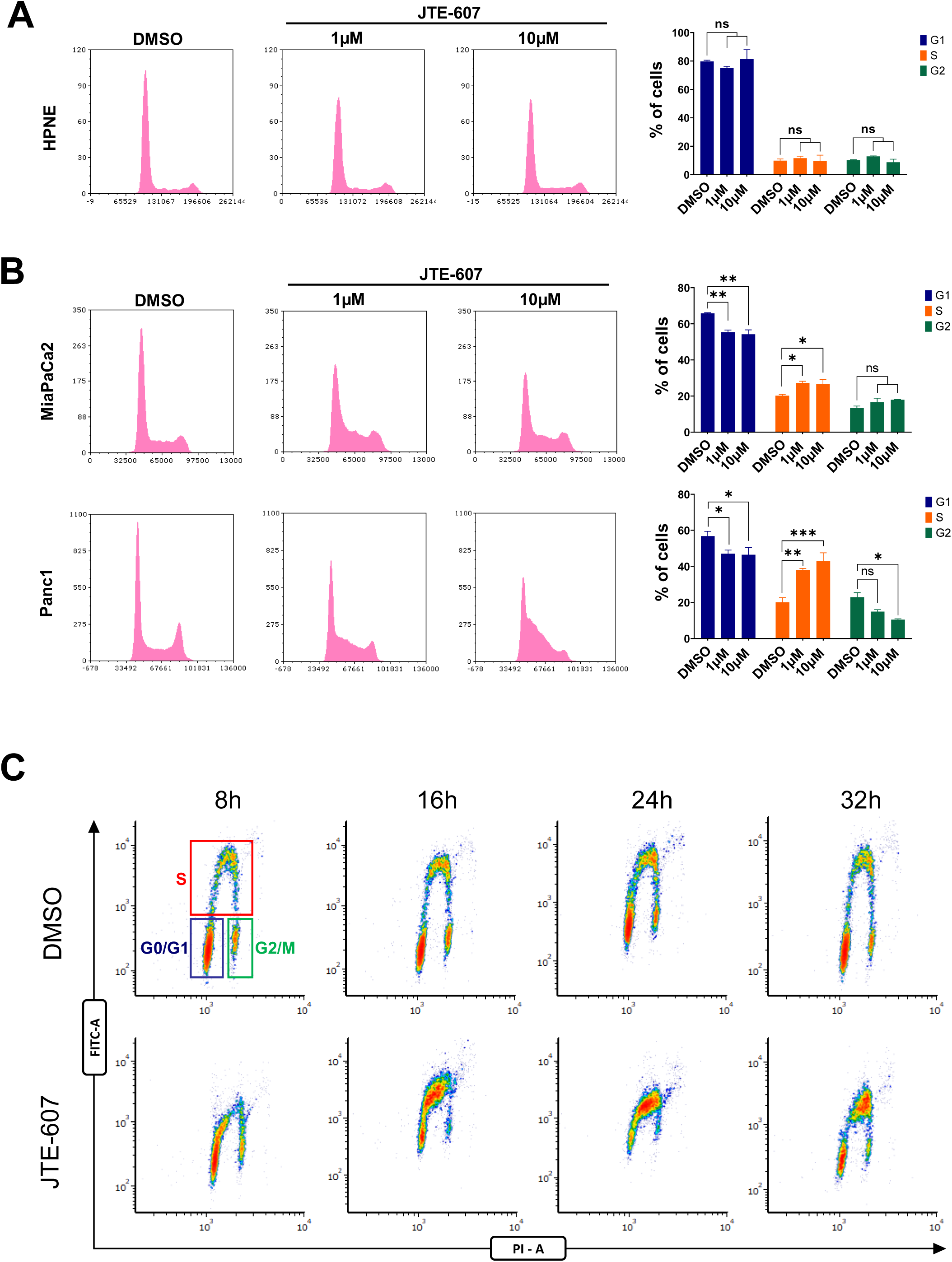
JTE-607 induces S-phase arrest. **A and B,** Cell cycle distribution and quantification of Panc1, MiaPaCa2 and HPNE cell lines. **\***, *P* < 0.05, **\*\***, *P* < 0.001, **\*\*\***, *P* < 0.0001, 2way ANOVA with Dunnett’s multiple comparisons test. **C,** BrdU incorporation assay showing cell cycle population upon JTE-607 treatment.

## Discussion

Dysregulation of gene expression is a fundamental driver of cancer (45). This dysregulation can be driven by non-mutational epigenetic reprogramming, a mechanism that is now recognized as a hallmark of cancer (46). Emerging evidence has implicated dysregulation of one such non-mutational gene regulatory process, APA, in the pathogenesis of cancer (23,47–49). Pan-cancer analyses have revealed global changes in APA across the cancer landscape, and mechanistic studies have characterized how these alterations promote oncogenesis (23,50–52). Recently, we reported the first large-scale, single cancer study of APA and discovered widespread alterations in 3’-UTR length across the PDAC landscape. Importantly, many of these APA alterations were associated with expression changes in growth-promoting genes, highlighting the importance of APA in driving PDAC pathogenesis (16). Therefore, we hypothesized that therapeutically targeting APA in PDAC would alleviate this global repatterning of 3’-UTR usage, normalizing the expression of oncogenes and tumor suppressors, and attenuating tumor growth. To directly test this hypothesis, we focused on CPSF3, an enzymatic component of the APA machinery that catalyzes the endonucleolytic cleavage of the pre-mRNA. While CPSF3 has known roles in the regulation of APA and histone mRNA processing, our study defines the first roles of CPSF3 activity in an epithelial cancer, with implications for therapeutic intervention in intractable pancreatic cancer. We demonstrate that high *CPSF3* expression is a predictor of poor patient outcome and uncover the requirement for *CPSF3* in PDAC cell proliferation *in vitro* and tumor growth *in vivo*. We characterize the global APA alterations driven by loss of *CPSF3*, revealing the dysregulation of numerous tumor suppressors and oncogenes. We then determine the direct connection between a CPSF3-mediated APA event in the tumor suppressor *FHL1* and PDAC cell proliferation. Finally, we reveal dysregulation of histone processing downstream of CPSF3 inhibition, regulating cell cycle progression. These results demonstrate the potential for targeting CPSF3 as a novel therapeutic approach in PDAC.

Our study has several clinical implications. First, we show that *CPSF3* expression is dysregulated in PDAC and high expression correlates with poor prognosis. This is consistent with similar findings across the cancer landscape, where *CPSF3* has been reported to be a predictor of unfavorable prognosis in lung and liver cancers (53, 54). While several studies have experimentally manipulated various APA factors and determined the phenotypic impacts, little is known about the function of CPSF3 in disease, particularly cancer. This is noteworthy for several reasons. First, CPSF3 is the enzymatic component of the APA machinery, and is thus a potentially druggable target. Second, despite acting in the same complex, knockdown of specific APA factors can have opposing impacts on APA and cellular phenotypes (50,51,55–58). Therefore, understanding the role of CPSF3 specifically in PDAC cell proliferation is critical for elucidating its potential as a novel therapeutic target. Recently, homozygosity in *CPSF3* missense variants was found to cause intellectual disability and embryonic lethality in humans. However, these phenotypes were completely absent in the heterozygous carriers (59). In cancer cell line models, *CPSF3* is essential for cell proliferation when knocked out completely by CRISPR; however, *CPSF3* is not an essential gene upon shRNA-mediated partial knockdown (www.depmap.org). This suggests that pharmacological targeting of such an essential gene may be biologically feasible. In support of this hypothesis, we show that knockdown of *CPSF3* blocks PDAC cell proliferation and tumor growth, and that the efficiency of knockdown is a determinant of phenotypic strength. Furthermore, CPSF3 inhibition does not impair cell cycle progression or proliferation of non-transformed pancreatic epithelial cells, and the CPSF3 inhibitor JTE-607 is non-toxic in humans. Therefore, inhibition of CPSF3 may preferentially target transformed cells.

Genetic manipulation of APA factors has been shown to alter APA patterns, dysregulate gene and protein expression and drive cancer phenotypes (50,51,63,64,52,53,55–57,60–62). While several recent reports have linked *CPSF3* loss to defects in tumor cell growth, no study has mechanistically connected *CPSF3* to global dysregulation of APA, gene expression and cell phenotype. We and others have previously shown the direct effect of cancer-associated APA alterations on gene expression and cancer cell proliferation and tumor growth (16,23,47,48). Adding to that body of literature, we now demonstrate that *CPSF3* knockdown alters 3’-UTR length of multiple tumor suppressors and oncogenes, including *FHL1, CMTM3, SMAD6 and MAP2K6*. To understand if these gene expression changes are responsible for driving *CPSF3* knockdown-mediated phenotypes, we determined the requirement for FHL1 expression in mediating cell proliferation. *FHL1* possesses tumor suppressing activity in different cancer types but has not been investigated in the context of PDAC (29–32). We find that silencing of *FHL1*, in the context of *CPSF3* knockdown, sufficiently rescues the defect in cell proliferation. Therefore, we uncover a novel mechanism by which CPSF3 loss attenuates PDAC cell growth. We also provide the first global view of the contribution of CPSF3 activity to PDAC gene expression, revealing widespread control of genes implicated in cell growth. While dysregulation of FHL1 is at least partially responsible for the *CPSF3* knockdown cell proliferation phenotype, we propose that *CPSF3* loss dysregulates the expression of a suite of cell growth genes through altering 3’-UTR length, thereby attenuating PDAC cell growth.

Recently, two groups independently demonstrated that CPSF3 is the target of the small molecule JTE-607 (18, 19). JTE-607 was first identified over 20 years ago as a cytokine synthesis inhibitor; however, the direct molecular target remained elusive. Despite the lack of a defined mechanism, JTE-607 was tested in a Phase I dose-escalation trial in healthy human volunteers, with no serious adverse effects (22). Therefore, despite inhibiting an essential enzyme responsible for processing >70% of polyadenylated mRNAs, JTE-607 is not uniformly toxic in humans. This property, coupled with our data demonstrating JTE-607’s anti-proliferative effects on cancer cells, supports the contention that targeting CPSF3 is a feasible prospect in PDAC. In humans, endotoxin-induced production of C-reactive protein, IL-10 and IL-1ra was inhibited by JTE-607 (22). In animal models, JTE-607 inhibited the production of proinflammatory cytokines, prevented endotoxin shock and attenuated artificially induced lung and heart injury (65–67). JTE-607 has also been used in models of acute myeloid leukemia (AML) and Ewing sarcoma and showed growth inhibitory activity both *in vitro* and *in vivo* (xenograft models) (19–21). However, these studies were limited to leukemia and sarcoma models, with no efficacy shown for epithelial-derived tumors. Therefore, the potential for CPSF3 as a therapeutic target in adenocarcinoma was an open question. Now, we show that JTE-607 preferentially blocks proliferation of PDAC cell lines, sparing non-transformed cell lines, including epithelial cells and fibroblasts. The mechanisms underlying this difference in sensitivity are currently unknown, but may relate to variability in basal proliferation rate. Finally, even though JTE-607 was first described as an inhibitor of cytokine synthesis, our RNA-seq analysis did not show an enrichment of such pathways. One possible explanation is that JTE-607 action is cell type dependent. Many of the studies assessing cytokine levels were performed using measurements from blood, and therefore the cell type responsible for the changes in cytokine secretion is unknown. It is possible that the effect of JTE-607 on proliferating epithelial cells is distinct from its effect on cells within the circulation, many of which are non-proliferative when terminally differentiated. The effects of JTE-607 in different cellular contexts and cell states warrants further investigation.

A previous report demonstrated that JTE-607 attenuates cell proliferation in AML and Ewing sarcoma through increasing R-loop formation and downregulating the expression of DNA damage response genes. R-loops are DNA:RNA hybrids that form as a result of aberrant transcription, a characteristic of cancers with genetic rearrangements such as AML and Ewing sarcoma (68, 69). Of note, R-loops increase in models with mRNA cleavage and polyadenylation defects (70), suggesting that sensitivity of AML and Ewing sarcoma to JTE-607 may be a consequence of high basal levels of R-loops, which eventually accumulate leading to DNA damage and genomic instability. In our study, gene set enrichment analysis did not reveal changes in DNA damage response pathways upon *CPSF3* knockdown or inhibition in PDAC cells. Therefore, we propose that CPSF3 regulates cell proliferation through distinct mechanisms in AML and Ewing sarcoma relative to PDAC. In PDAC cells, we find that JTE-607 impairs processing of proliferation-dependent histone mRNAs. This is consistent with the role of CPSF3 in the HCC (17,38,71–73). Defects in the HCC have been shown to reduce the availability of replication-dependent histones (17,40,74,75). However, prior to now, no studies have described the effect of CPSF3 inhibition on HCC activity. Depletion of many HCC genes led to an accumulation of histone read-through transcripts in the nucleus (37, 39). Similarly, we find extensive transcript read-through in proliferation-dependent histone mRNAs, but not proliferation-independent histone mRNAs upon JTE-607 treatment. Histone read-through transcripts accumulate in the nucleus, thus failing to be exported into the cytoplasm and translated into protein. In accordance with this model, we find that JTE-607 depletes core histone protein levels in PDAC cells. Limited histone supplies destabilize chromatin integrity through disruption of nucleosome assembly (43). We find that JTE-607 destabilizes chromatin stability, as demonstrated by increased sensitivity to MNase digestion, and derepression of heterochromatin-mediated gene expression silencing. These findings reveal a novel mechanism of JTE-607 activity: dysregulation of proliferation-dependent histone mRNA processing.

Expression of replication-dependent histones increases ∼30-50 fold during DNA synthesis (76, 77). The life cycle of these core histone genes starts late in G1 through mid S phase of the cell cycle and degradation occurs at late S phase (36, 78). Silencing of the HCC core component *FLASH* induces S phase arrest in HeLa cells (79). We find that JTE-607 arrests cells in the S phase of the cell cycle, with cells slowly cycling through early-mid S phase but failing to progress through late S phase. This is consistent with a previous study where depletion of the histone chaperone ASF1, an important gene for histone deposition during DNA replication, disrupts progression through mid to late S-phase (80). In addition, knockdown of CSTF2, a gene with dual functions in APA and histone pre-mRNA processing, delays progression through S phase, but its expression is highly dependent on cell cycle stage (39). The same study showed that CPSF3 expression is not cell cycle regulated, suggesting that the histone phenotype we observe is driven by CPSF3 loss and not merely a consequence of cell cycle arrest. Our findings strongly suggest that JTE-607 mediates its growth attenuating phenotype by reducing histone supplies during S phase, thereby blocking cell cycle progression.

This newly discovered mechanism of JTE-607 represents a potential window for new combination therapy. One possibility is that JTE-607-mediated cell cycle arrest may promote synergism with cell cycle check-point inhibitors. For instance, the chromatin remodeling histone deacetylase (HDAC) inhibitors have shown synergistic effect when combined with checkpoint kinase 1 (Chk1) inhibitors in lung cancer models (81). Histone disruption by JTE-607 may also promote synergism with chromatin modifying drugs. For example, CBL0137 has shown synergistic effect when combined with HDAC inhibitors by exacerbating chromatin destabilization (44). These discoveries may improve the efficacy of approved chromatin remodeling agents and suggest a path forward for use of JTE-607 in the clinic.

The mechanistic differences underlying the CPSF3 knockdown and inhibition phenotypes raises several important questions. As CPSF3 is an integral subunit of both the APA and histone processing complexes, *CPSF3* knockdown may disrupt proper recruitment of other complex components. This dysregulation of complex formation may alter complex function in different ways than inhibition of CPSF3 activity, resulting in divergence in APA patterns and gene expression alterations. Furthermore, the process of generating stable cells for long-term *CPSF3* knockdown can result in upregulation of compensatory mechanisms, allowing cell growth in the absence of an essential gene. These mechanisms will not be accounted for upon pharmacological CPSF3 inhibition, again resulting in differences in APA patterns and gene expression alterations. However, it should be noted that both CPSF3 knockdown and inhibition led to decreased cell proliferation and global dysregulation of APA, including that of known tumor suppressors and oncogenes. Therefore, our results support the development of CPSF3 targeting agents, including those that can specifically degrade CPSF3.

In conclusion, our study has revealed the role of CPSF3 in pancreatic cancer and uncovered new mechanisms by which CPSF3 mediates cell proliferation. CPSF3 knockdown or inhibition induces APA changes that alter the expression of known tumor suppressors and oncogenes. CPSF3 inhibition disrupts the processing of proliferation-dependent histones, destabilizing chromatin structure and inhibiting cell cycle progression. Our findings reveal novel insight into how CPSF3 inhibition blocks cell proliferation and provides a new therapeutic target in pancreatic cancer.

## Authors’ Disclosures

The authors declare no potential conflicts of interest

## Authors’ Contributions

**A.A. Alahmari:** Conceptualization, methodology, investigation, data analysis, writing–original draft, writing–review and editing. **A.H. Chaubey:** Investigation. **A.A. Tisdale:** Investigation. **C.D. Schwarz:** Investigation. **A.C. Cornwell:** Investigation. **K.E. Maraszek:** Investigation. **E.J. Paterson:** Investigation. **M. Kim:** Investigation. **S. Venkat:** Investigation, bioinformatics analysis. **E.C. Gomez:** Bioinformatics analysis. **J. Wang:** Bioinformatics analysis. **K.V. Gurova:** Resources. **M.E. Feigin:** Conceptualization, supervision, funding acquisition, project administration, writing–review and editing.

## Acknowledgements

This work was supported by National Cancer Institute (National Institutes of Health) grants P30 CA016056 and R25 CA181003, an award from the Roswell Park Alliance Foundation to M.E.F., and a scholarship and support to A.A.A. from Prince Sattam bin Abdulaziz University in Saudi Arabia, the Saudi Arabian Cultural Mission in the USA, and the Office of International Collaborations in Oncology at Roswell Park Comprehensive Cancer Center. M.E.F. is supported by a Research Scholar Grant, RSG-21-014-01 - RMC, from the American Cancer Society. We thank the Roswell Park Genomics and Small Molecule Screening Shared Resources for their assistance. We thank Safina Alfyia for her help with the MNase experiment and Brian Buckley for assistance in imaging HeLa-TI cells. We thank the members of the Abel Laboratory for their insightful comments and support.

## Notes

https://github.com/feiginlab/CPSF3_PDAC

### Competing Interest Statement

The authors have declared no competing interest.

